# Gastric Cancer Based on Transcription Factors

**DOI:** 10.1101/2025.09.07.674761

**Authors:** Aline Gonçalves da Costa, Kauê Sant’Ana Pereira Guimarães, Ronald Matheus da Silva Mourão, Juliana Barreto Albuquerque Pinto, Jéssica Manoelli Costa da Silva, Diego Pereira, Valéria Cristiane Santos da Silva, Samia Demachki, Williams Fernandes Barra, Samir Mansour Casseb, Fabiano Cordeiro Moreira, Rommel Mario Rodriguez Burbano, Paulo Pimentel de Assumpção

**Author notes:** Address of corresponding author: Aline Gonçalves da Costa.

## Abstract

**Introduction:** Gastric adenocarcinoma (GA) is responsible for thousands of deaths annually and significantly affects patients’ quality of life. To increase knowledge about this disease, several studies have focused on the transcriptional control mechanism by analyzing the role of transcription factors (TFs) in GA.

**Methods:** This study identified differential expression (DE) of TFs in GA samples from 74 patients using NGS, biostatistical analysis in R, and comparison with 68 normal tissue samples in the genetic databases of the National Center for Biotechnology Information (NCBI).

**Results:** 1,564 human transcription factors were DE. Of these, 87 were selected using the established cutoff value of AUC ≥ 0.95, and 25 were analyzed for their influence on GA carcinogenesis. Based on their role in biological processes, nine TFs were found to be underexpressed due to their growth suppression and cell differentiation properties, notably the TFs IRF3 and CXXC1. Sixteen TFs were overexpressed, associated with drug resistance, disease promotion, metastasis, and poor prognosis in GA, such as AHR, NCOA3, RRB1, and STAT3, respectively.

**Conclusion:** This study integrated systematic bioinformatics and NGS, which revealed novel TFs with important oncogenic and suppressive functions in GA. Therefore, it provides new insights into genes with potential biomarkers for prognostic prediction in GA, which may offer important implications for clinical practice and may effectively target treatment, monitoring, or disease recurrence.

## INTRODUCTION

Gene regulation is the process by which cells control which genes, present in the genome, will be “turned on” or “turned off” at various points of cellular differentiation. Although almost all eukaryotic cells have exactly the same DNA, different gene expression patterns allow different cell types to possess diverse sets of proteins with tissue-specific functions. However, genetic alterations can affect gene expression and predispose to tumor development^(1)^.

Cellular malignancy is an example of deregulation of gene expression, which is a consequence of the accumulation of mutations in DNA, which, in turn, causes biological and functional damage to cells^(2)^. Among these is gastric cancer, one of the most common human cancers, representing a significant public health problem worldwide with 90% of cases classified as gastric adenocarcinoma (GA)^(3)^.

In recent years, the scientific community worldwide has made a significant effort to interpret the wealth of information related to gene regulation obtained by DNA sequencers and its association with cancer development and progression. To this end, studies have evaluated the transcriptional control of DNA, carried out by transcription factors (TFs), discovered in 1961^(4)^.

From this perspective, TFs are related to prognoses in GA^(4)^, assisting in the stratification of high-risk patients, as well as in guiding treatment^(5,6)^. Among them, there are those that express proteins involved in cell cycle control, DNA damage repair and apoptosis induction, as well as those that regulate the process of cell invasion, adhesion, epithelial-mesenchymal transition (EMT) and oncogene expression^(7,8)^.

Based on the above, the identification and descrption of the main TFs involved in gastric carcinogenesis is the central point of the present study.

## MATERIAL AND METHODS

### Ethics statement

This study was approved by the Ethics and Research Committee of João de Barros Barreto University Hospital (approval number: 47580121.9.0000.5634) and was conducted in accordance with the principles outlined in the Declaration of Helsinki. Participant recruitment and sample collection were carried out between July 2, 2022, and July 6, 2023. Before enrollment, all participants received detailed information about the study’s objectives, potential benefits, risks, and possible harms, ensuring a thorough understanding of the research. Written informed consent was voluntarily obtained from all participants prior to their inclusion in the study.

### Sample Collection

In addition, data were obtained from the PRJNA1054173 project, available in Bioproject NCBI (https://www.ncbi.nlm.nih.gov/bioproject/?term=PRJNA1054173), which contains the transcriptome of gastric tissues from patients without cancer.

A descriptive study of TFs identified in GA samples was carried out. For this purpose, 142 samples were used, 74 of which were from GA tissue and 68 from normal tissue, identified on platforms that gather reference genes for biomolecular research.

### Extraction and quantification of total RNA

To obtain total RNA, tumor tissue macerates were used, performed using the TRIzol Reagent RNA extraction method (*Thermo Fisher Scientific*), following the manufacturer’s instructions. Then, the flurimetric method was performed to determine the integrity and concentration of RNA, using the Qubit 4 Fluorometer equipment (*Thermo Fisher Scientific*). The established criteria for RIN (RNA integrity numbers) were ≥ 5.

### Genomic library construction and sequencing

The genomic library was prepared and sequenced using the methodology described in the TruSeq Stranded Total RNA Library Prep Kit with Ribo-Zero Gold (Illumina, US). After library completion, the samples underwent an integrity assessment of the generated DNA fragments, performed on the 2200 TapeStation System (Agilent Technologies AG, Basel, Switzerland).

The genome assembly sequence was performed in pair-end on the NextSeq® 500 platform (Illumina®, US), using the NextSeq® 500 MID Output V2 kit - 150 cycles (Illumina®), according to the manufacturer’s instructions.

### Quality control of sequencing data

After sequencing, the RNA-Seq reads were subjected to quality control. Initially, the raw sequences were quality assessed using FastQC v0.11.9. Subsequently, Trimmomatic v0.39 was used to remove adapters and low-quality reads. The files containing the filtered sequences were re-evaluated in FastQC.

### Alignment of samples with the human genome

Human transcript reads were characterized by alignment and quantification with Salmon v1.5.2, using the coding transcripts in hg v38 (www.ensembl.org) as a reference index, in addition to using GENCODE v.42 (www.gencodegenes.org) as annotation. Reads were imported from Salmon into RStudio with the Tximport v3.14.0 package.

### Multivariate Analysis

#### Differential Expression (DE) Analysis of Human Genes

After classification and alignment, a differential gene expression analysis was performed between GA tumor samples and normal tissue, using the DESeq2 package^(9)^. Genes considered differentially expressed (DE) were those whose expression difference met the criteria: I) |Log2(Fold-Change)| > 1; and II) adjusted p-value < 0.05.

The results of differential gene expression were visualized using the ggplot2 v3.5.1^(10)^ and complexHeatmap v2.14.0 packages.

#### Receiver Operating Characteristic Analysis

To determine the specificity and sensitivity of differentially expressed (DE) genes, the area under the curve (AUC) of the receiver operating characteristic (ROC) was calculated using the pROC v1.18 package^(11)^. Genes with AUC > 0.95 were selected.

#### Functional enrichment analyses of genes

DE genes with AUC > 0.95 were enriched using Gene Ontology for terms related to biological processes to better understand their biological and functional relevance. The Cluster Profile v4.8.3 package was used for this analysis.

#### Gene Expression Analysis

To evaluate the expression of TFs, the identified samples were selected and submitted to the nf-core/rnaseq v3.14 pipeline. Reference genomes available in the *National Center for Biotechnology Information* (NCBI) database were used as indexes.

#### Analysis of TF expression in GA tissue compared to normal tissue

To compare the expression of TFs in tissue with GA and normal tissue, PCA (Principal Component Analysis) was performed, which is frequently used to analyze genomic data and identify significant variations between different samples.

## RESULTS

### Differential Gene Expression Analysis

Our results identified 1,564 human transcription factors differentially expressed between the samples, according to the criteria established for DE genes. Of these, 87 were selected using the established cutoff value for AUC and 25 (FIGURE 1) were analyzed for their influence on GA carcinogenesis.

**FIGURE 1.**
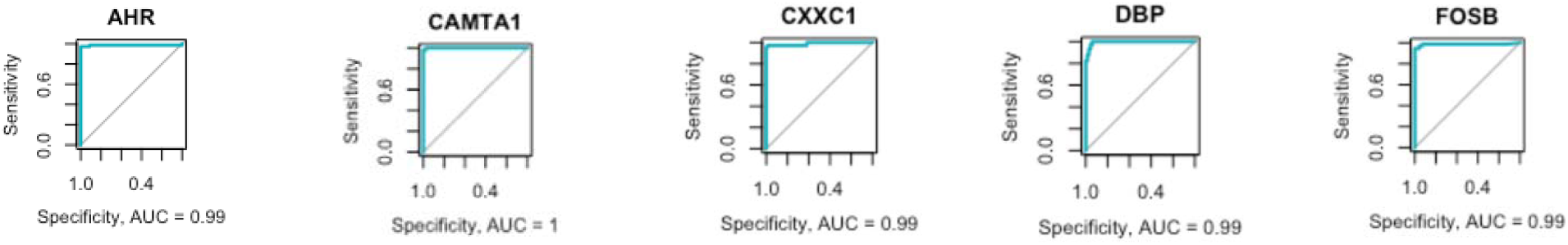
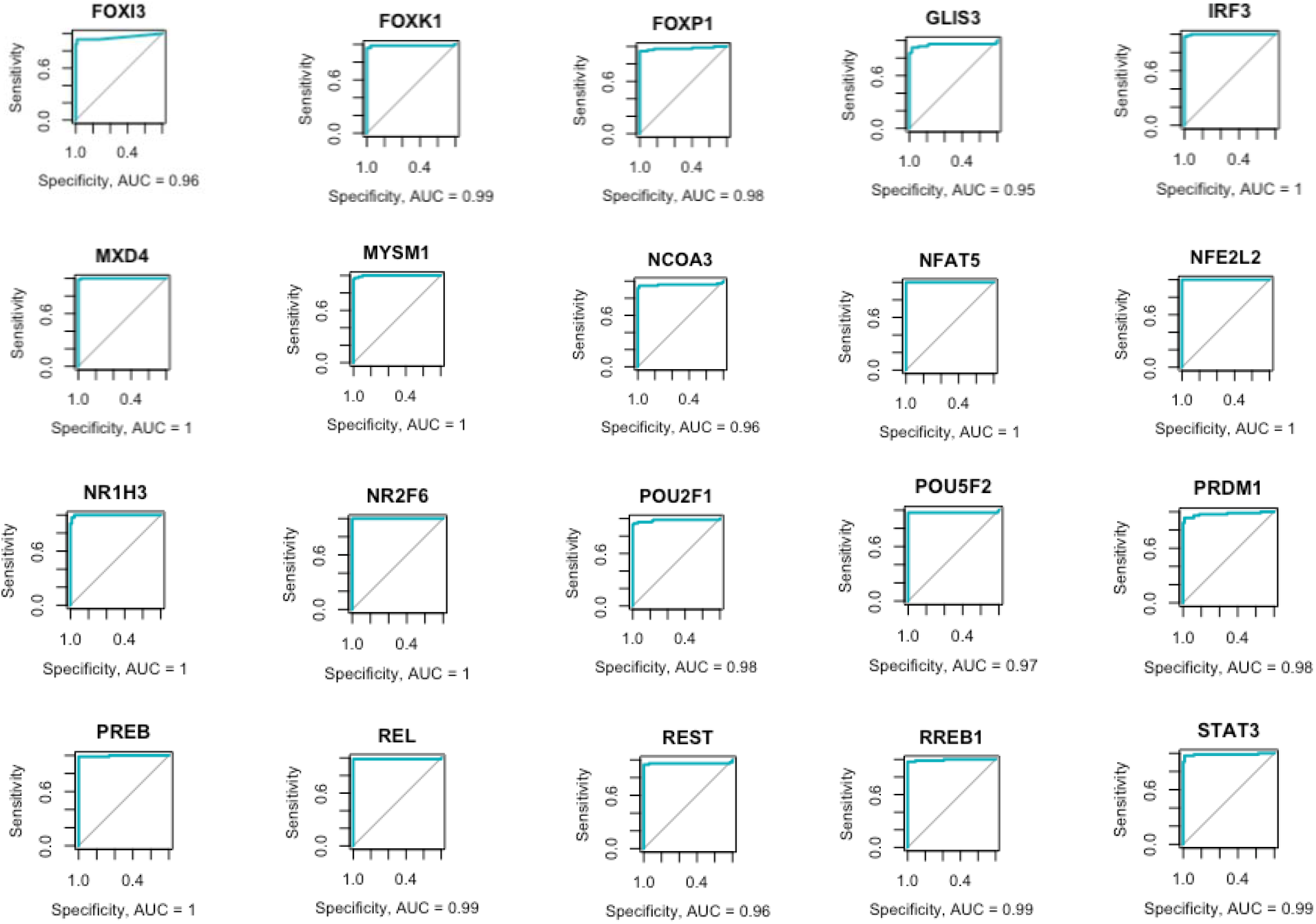
ROC curve graphs representing the 25 DE transcription factors analyzed in gastric adenocarcinoma.

Then, the samples were classified as hypoexpressed and hyperexpressed (FIGURE 2), considering the differential gene expression profile between tissues with GA and normal tissue.

**FIGURE 2.**
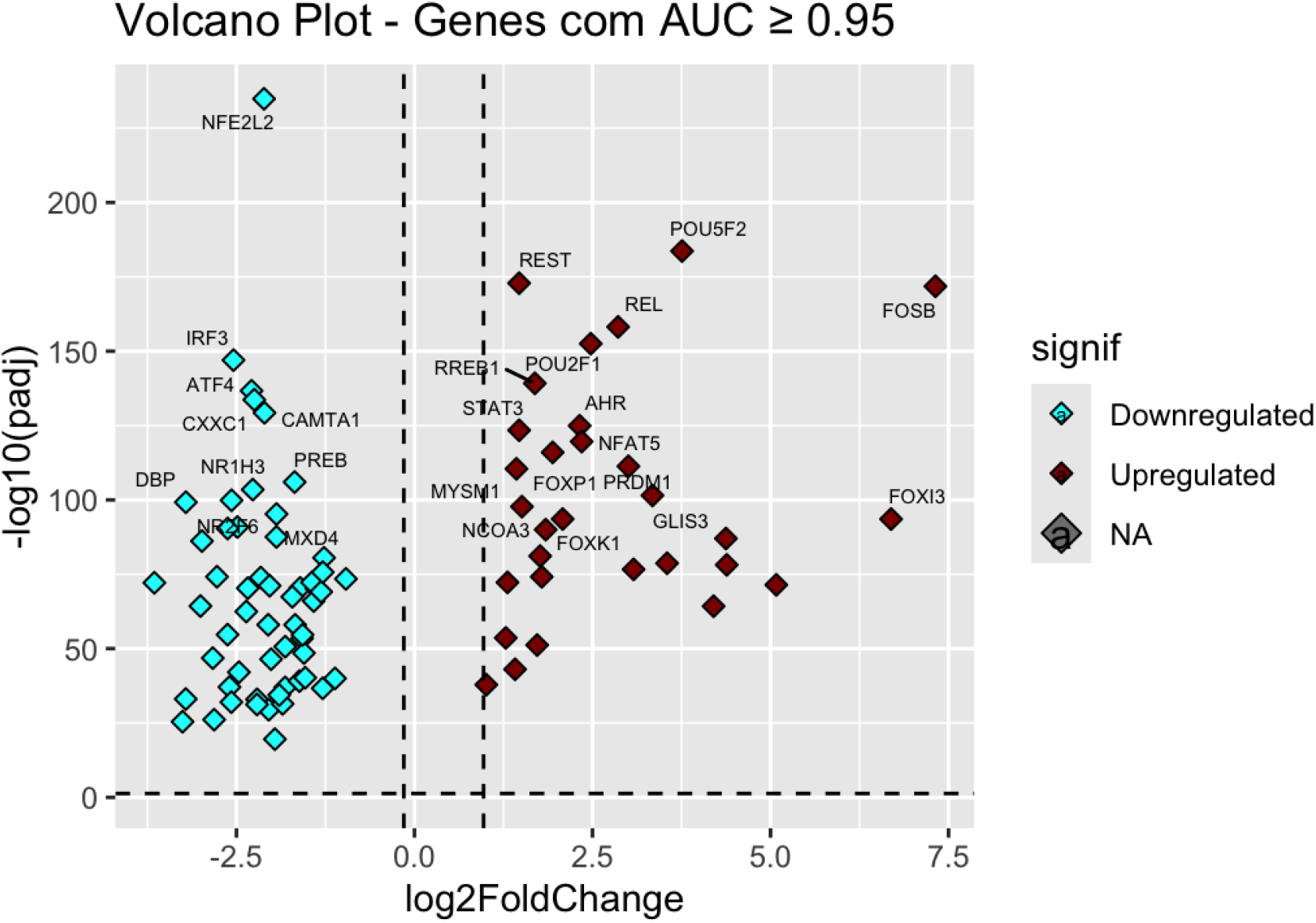
Volcano plot showing DE genes among samples that showed transcription factors with AUC ≥ 0.95. In green are the hypoexpressed TFs and in red the hyperexpressed TFs.

To better visualize the gene expression patterns between the GA and normal tissue samples, an unsupervised hierarchical clustering analysis was conducted. This revealed that the GA samples exhibited a distinct gene expression pattern from that of the normal tissue samples, which can be seen in the formation of clusters, demonstrating that certain groups of TFs exhibit more intense upregulation in GA samples than in normal samples. Furthermore, the *heatmap* (FIGURE 3) demonstrates that the samples were grouped according to their similar expression patterns.

**FIGURE 3.**
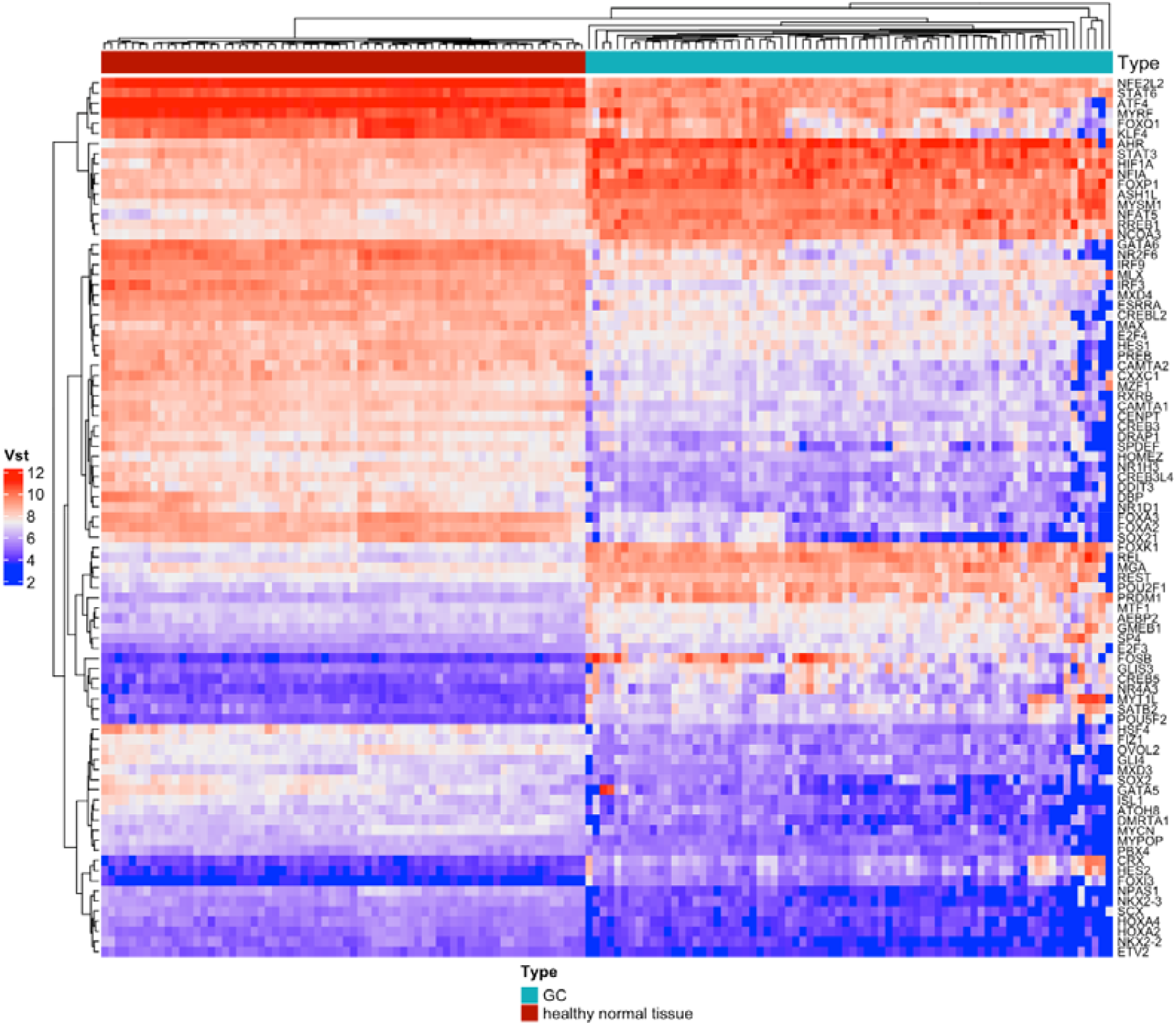
Heatmap of the expression patterns of DE TFs with AUC ≥ 0.95.

### Variation of the expression of DE TFs in GA in relation to normal tissue

After analyzing the expression of TFs in the samples, the DE TFs in the GA tissues were compared with the DE TFs in the normal tissues indexed in the databases. It was possible to observe the formation of two larger clusters that were significantly expressed and differentiated between the samples (FIGURE 4).

**FIGURE 4.**
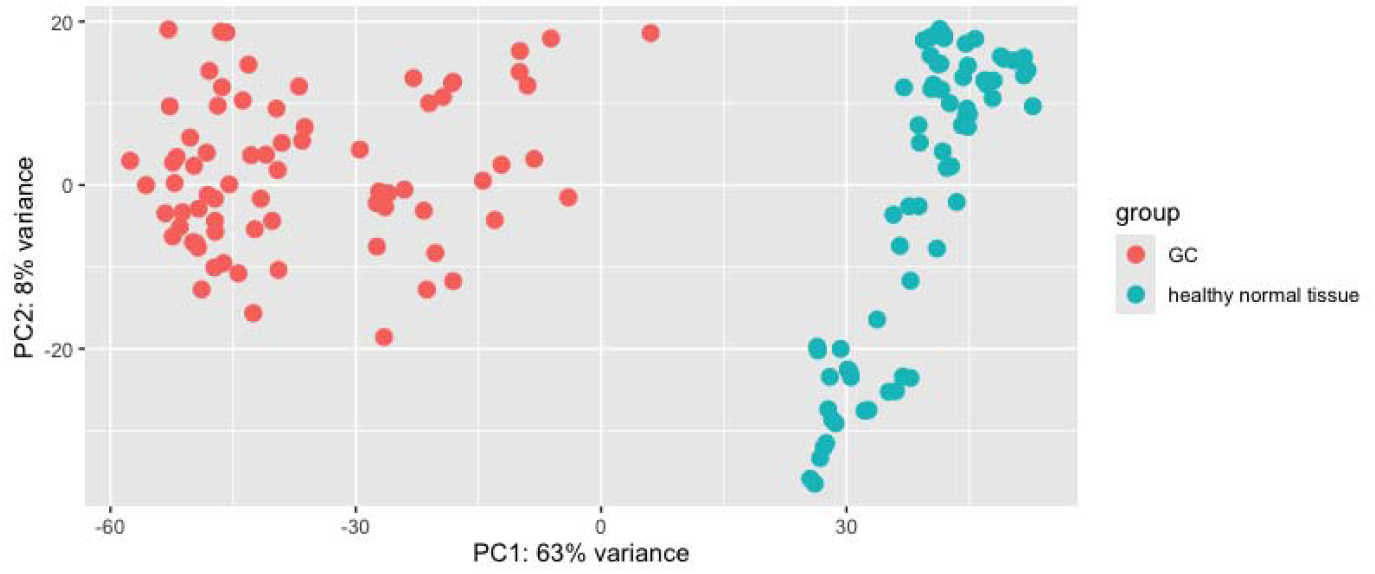
Variation between DE TFs with AUC ≥ 0.95 expressed in GA and normal tissue samples from genetic databases.

## DISCUSSION

The identification of TFs in GA samples has considerable clinical relevance because, in addition to specific molecular and histological characteristics, patients with the presence of these genes can be targeted for distinct and more precise therapeutic approaches in the future. Therefore, we sought to analyze the implications of the main expressed TFs in gastric carcinogenesis.

In the literature, there are several studies that have already been conducted on the expression of TFs in GA, whose positive regulation correlates with better survival; with suppression of proliferation, cell migration and metastasis; with inhibition of growth and angiogenesis and with reduction of tumors and cells resistant to chemotherapy. In contrast, most TFs identified in the studies promote cell proliferation, cancer progression, invasion and migration of GA cells, facilitate glycolysis, as well as metastasis, apoptosis, drug resistance, angiogenesis, stemness and metabolism of gastric cancer cells, in addition to being associated with worse survival in patients with GA^(6,12,13,14)^.

Therefore, our study analyzes the identified TFs, relating them to gastric carcinogenesis and whether, in the near future, they can be considered biomarkers of GA. Thus, the TFs were classified into two groups: a) underexpressed TFs and b) overexpressed TFs, according to their activity and relationship to the oncogenic process.

### A) Underexpressed TFs

The first to be described is the TF CAMTA1 (calmodulin-binding transcription activator 1), found to be underexpressed in GA samples. In the literature, this TF plays an important role in delaying cell proliferation and tumor growth in neuroblastomas^(15,16)^. Therefore, it is understood that CAMTA1 and its variants can also influence the tumor microenvironment in GA, which may explain its low expression in the samples, indicating possible progression of GA in the analyzed tissues.

Following the analysis of underexpressed TFs, the TFs IRF3 and CXXC1 are reported in the literature to play an important role in regulating various genes or signaling pathways to prevent GA progression or inhibit the proliferative potential of cancer cells. For example, when IRF3 is upregulated and associates with YAP and TEAD4, prognostic analysis of patient survival is possible^(17)^. Similarly, CXXC1 acts by interfering with the binding of proteins (ELK1 to the miR100HG promoter) that would promote cell proliferation, increasing the secretion of IFN-γ from CD3+ T cells and, consequently, inhibiting immune escape in GA cells^(18)^.

In the same way, nuclear receptors (NRs), such as the TF NR2F6, are involved in the regulation of immune and tumor cells, limiting cell growth through the infiltration of immune cells, mainly TCD8+, in a study with xenografts and mice for melanomas^(19)^. Furthermore, there are also studies in which FT PREB may act in the regulation of prolactin and its cognates, participating in the mechanisms of inhibition or expression of breast cancer^(20)^. In the studied samples, hypoexpression is found, indicating tumor invasiveness in the GA.

There are also TFs that act in the cis-regulatory region of RNA polymerase II, such as NR1H3, and those that regulate antioxidant response elements and ferroptosis induction, such as NFE2L2, despite their duality in cellular balance in lung cancer therapy^(21)^. However, there are no studies described in the literature on these TFs and their expression in GA, this being the first analysis performed.

The TF MXD4 was underexpressed in the GA samples. A similar correlation was found in the literature in a bioinformatics study with gastric cancer tissue. This TF and other genes in the MXD family may serve as biomarkers of poor prognosis or as potential targets for immune therapy, since they regulate endocytosis, the cell cycle, and apoptosis^(22)^. In addition, there are other studies associating MXD4 with the suppression of oncogenesis in leukemia^(23)^.

The TF DBP is underexpressed in the samples, and no descriptions linking it to GA were found. However, there is research suggesting that acetylation in promoter regions of this gene may play an important role in glucose energy metabolism, influencing adipose tissue in patients with type 2 diabetes, and that polymorphisms in specific DBP components are associated with cancer risk^(24,25)^.

Therefore, it is inferred that the 09 hypoexpressed TFs described above present growth and cell differentiation suppression properties, which support their attribution as tumor suppressor genes in GA.

### B) Overexpressed TFs

As previously mentioned, most TFs identified to date are related to drug resistance, promotion, metastasis, and poor prognosis. Among them is STAT3, which acts as a negative regulator of ferroptosis, leading to chemoresistance in GA, although research indicates a certain susceptibility of cancer cells in vitro to ferroptosis/oxytosis^(12,26)^. In our findings, TFs are overexpressed in the studied GA tissue samples, which may be related to a poor prognosis and cell migration^(27)^.

Overexpression of PRDM1, a tumor suppressor gene, has been described through its polymorphism in association with programmed cell death ligand 1 (PD-L1), influencing the size of hepatocellular carcinoma tumors^(28)^. In the GA tissue samples from this study, TFs are overexpressed, which may indicate an active immune response against tumor cells, since PRDM1 overexpression increases PD-L1 levels, favoring the antitumor immune response^(29)^. Therefore, the findings of this study indicate PRDM1 as a possible predictive biomarker for responses of patients with GA to immunotherapy.

The TF NFAT5 (nuclear factor of activated T cells 5), also overexpressed, is linked to malignancy and increased migration of esophageal cancer cells, through regulation of the TLR4/MyD88 signaling pathway, in addition to being associated with poor prognosis in pancreatic adenocarcinomas^(30,31)^. In contrast, the TF MYMS1, a metalloprotease, which is overexpressed in the studied GA samples, plays a suppressive potential and is associated with a favorable prognosis in cases of colorectal cancer, in addition to suppressing oncogenesis through the inhibition of PI3K/AKT signaling^(32)^, which can be evaluated a posteriori to confirm these hypotheses also for GA.

In addition, the TFs FOXK1 and FOXP1 demonstrated upregulation in GA tissue samples. In the literature, an association with GA was found only of FOXK1, in which the TF is described as an important regulator of cell proliferation, quiescence, and differentiation in stem cell populations and malignancy for GA^(33)^. FOXP1, on the other hand, functions as an oncogene in ovarian cancer cells, promoting cancer stem cell-like characteristics^(34)^. Overall, Foxo factors have been shown to inhibit apoptosis, regulate PI3K signaling, and unwrap chromatin to expose DNA-binding motifs for gene expression regulation^(35)^.

The TF AHR, an aryl hydrocarbon receptor, is frequently reported in tumorigenesis and may act as a positive or negative regulator of carcinogenesis. Studies show that sustained AHR activation induces the expression of genes associated with resistance to BRAF inhibitors in the treatment of metastatic melanoma, as well as mediates the activation of PI3K/Akt and MEK/ERK signaling via Src kinase, inducing cellular resistance. This TF plays an important role in gastric cancer, since the use of PKC (Protein Kinase C) inhibitors can control the epigenetic regulation of AHR^(36)^. This gene was also found to be overexpressed in GA samples, requiring prognostic validation.

On the other hand, NCOA3, also known as steroid receptor coactivator-3 (SRC-3), is described as a genuine oncogene and its overexpression indicates that it is sufficient to initiate the tumorigenesis process in several types of cancer, including breast cancer. Furthermore, it is observed that this TF integrates various signaling pathways, such as nuclear receptor, HER2/neu, IGF/AKT and NFκB, which are frequently involved in cell proliferation, survival and migration^(37)^. Thus, it is inferred that the overexpression of NCOA3 in the study results may be involved in the promotion or progression of GA carcinogenesis.

Following the analyses, overexpression of GLIS3 was identified in the samples. Upregulation of this TF was associated with poor prognosis and invasiveness in studies with GA cell lines, through regulation of the TGF-β1/TGFβR1/Smad1/5 pathway, which facilitates the malignant phenotype^(38)^. Other findings also demonstrated that GLIS3 promotes GA progression through immune infiltration, activation of microRNAs, such as miR-1343-3p, and vimentin phosphorylation^(39,40)^. All these findings support the hypothesis of this TF as a potential biomarker or therapeutic target for GA patients.

The TF FOXI3 gene, also overexpressed, has not yet been described in association with GA, although when dysregulated, it can cause genetic disorders. Literature findings have shown that pathogenic variants of this gene are linked to craniofacial microsomia, with autosomal dominant inheritance with reduced penetrance or autosomal recessive inheritance^(41,42)^. Therefore, phenotypic risks depend on the identified variants and their degree of penetrance. To extrapolate these hypotheses to GA, further studies on its possible influence on gastric carcinogenesis are needed.

Likewise, FT FOSB has been linked to the promotion of growth and metastasis in liver cancer, when associated with the MAFG-MM-7 axis, by inducing increased expression of the proteolytic enzyme MMP-7 in endothelial cells^(43)^. Furthermore, there is also a study in which this TF is directly linked to the evolution of non-small cell lung cancer; however, its role in tumor biology is determined by the genetic background of TP53 and its interactions, which lead to the selective transcriptional activation of distinct target genes that may interfere with the progression and prognosis of the disease^(44)^. However, there is no description in the literature of any study of FOSB hyperexpression in GA, as identified in the current findings.

The RREB1 oncogene has been described as highly expressed in GA, but its role in the tumor microenvironment is negatively related to the tumor suppressor protein p16, in addition to interfering with the immunological and T cell receptor microenvironments during neoadjuvant immunotherapy^(45,46)^. In the study results, TF is hyperexpressed in GA samples, which may indicate cell proliferation, due to an active immune response, tumor growth and lymphovascular invasiveness, caused by this gene^(47)^.

Similarly overexpressed, but with dual descriptions in the literature, is the TF REL. This TF has members, such as RelA and RelB, which are related to the development and maintenance of pancreatic ductal adenocarcinoma, through the downregulation of suppressor molecules (arginine 1) or by suppressing T cell function^(48)^. On the other hand, the c-Rel member is sometimes reported as a tumor suppressor, when it promotes immunosuppressive transcription in myeloid-derived suppressor cells - through a unique transcriptional complex called the c-Rel enhancer - and sometimes as an oncogene, when it promotes pro-inflammatory transcription in macrophages^(49,46)^. Thus, an important role of this TF in the immune response in cancer is observed.

Despite being highly expressed in the samples studied, no correlation was found between TF REST and GA. However, there are studies that link it to other types of cancer. Among them, one found a negative association between REST inhibition and the upregulation of the proteins CEMIP and MMP24, which play roles in metastasis and invasion in breast cancer^(50)^. There is another that associates REST regulation with the phenotypic control of neuroendocrine differentiation in small cell lung cancer^(51)^. Thus, the presence of this TF may enable prognostic evaluations in GA.

Finally, it is important to highlight that members of the POU domain gene family encode transcriptional regulatory molecules with important roles in the body. In vitro studies indicate a strong interaction between the POU class 2 TF domain 1 (POU2F1) and the ability to transcriptionally activate lncRNAs (TTC3-AS1 and LINC01564), promoting cell viability, migration, tumor progression, and invasion in GA^(52,53)^. Furthermore, POU2F1 overexpression, as in the present study, can also promote metastasis through downregulation of microRNAs in GAAG tissues^(54)^. However, no relationship was found between POU5F2 overexpression and AG or other cancer types.

## CONCLUSION

Therefore, the results discussed so far suggest a key role for DE TFs in both suppression and tumor development and progression in GA, through the up- or downregulation of other TFs, RNAs, or proteins. This study integrated systematic bioinformatics and NGS, which revealed novel TFs with important oncogenic and suppressive functions in GA. However, the incorporation of experimental methods is essential to corroborate the findings described. This study, therefore, provides new insights into genes with potential biomarkers for prognostic prediction in GA.

## Acknowledgment

The authors would like to thank the Oncology Research Center, the Human and Medical Genetics Laboratory, and the Anatomical Pathology Laboratory at João de Barros Barreto University Hospital (HUJBB – UFPA) for their invaluable technical and laboratory support. Our gratitude also goes to the High-Performance Computing Center (CCAD) at the Federal University of Pará for access to the Apollo 2000 cluster, which was crucial for our analyses.

## Data availability statement

The original contributions presented in the study are included in the article. Further inquiries can be directed to the corresponding author.

## REFERENCES

1 Reece JB, Urry LA, Cain ML, et al. Figure 18.6. Stages in gene expression that can be regulated in eukaryotic cells. Campbell Biology - 10th ed. San Francisco, CA: Pearson; 2011.

2 Griffits AJF, Doebley J, Peichel C, Wassarman DA, eds. Introdução à genética – 9th ed. Rio de Janeiro, RJ: Editora Guanabara Koogan; 2008.

3 Wild CP, Weiderpass E, Stewart BW. eds. World cancer report: cancer research for cancer prevention. Lyon (FRA): International Agency for Research on Cancer; 2020.

4 Zhou YY, Kang YT, Chen C, et al. Combination of TNM Staging and Pathway Based Risk Score Models in Patients with Gastric Cancer. J Cel Biochem. 2018;119(4):3608–3617.

5 Xu J, E C, Yao Y. et al. Matrix metalloproteinase expression and molecular interaction network analysis in gastric cancer. Oncology Letters. 2016;12(4):2403–2408.

6 Yang WJ, Zhao HP, Yu Y, et al. Updates on global epidemiology, risk and prognostic factors of gastric cancer. World J Gastroenterol. 2023; 29(16):2452–2468.

7 Zhao Z, Zhang C, Zhao Q. S100A9 as a novel diagnostic and prognostic biomarker in human gastric cancer. Scandinavian Journal of Gastroenterology. 2020;55(3):338–346.

8 Saito M, Kono K. Landscape of EBV-positive gastric cancer. Gastric Cancer. 2021;24(5):983–989.

9 Love MI, Huber W, Anders S. Moderated estimation of fold change and dispersion for RNA-seq data with DESeq2. Genome Biology. 2014;15(12):550.

10 Wickham H, Chang W, Henry L, et al. ggplot2: Elegant Graphics for Data Analysis. ggplot2 Web site. https://ggplot2.tidyverse.org. Accessed October 17, 2024.

11 Robin X, Turck N, Hainard A, et al. pROC: an open-source package for R and S+ to analyze and compare ROC curves. BMC Bioinformatics. 2011;12:77.

12 Wang T, Fahrmann GF, Lee H, et al. JAK/STAT3-Regulated fatty acid beta-oxidation is critical for breast cancer stem cell self-renewal and chemoresistance. Cell Metabol. 2018;27(1):136–150.

13 Cheng C.-W, Chang C.-C, Patria YN, et al. Sex hormone-binding globulin (SHBG) is a potential early diagnostic biomarker for gastric cancer. Cancer Medicine. 2018;7(1):64–74.

14 Li S, Yu W, Xie F, et al. Neoadjuvant therapy with imune checkpoint blockade, antiangiogenesis, and chemotherapy for locally advanced gastric cancer. Nature Communications. 2023;14(8):1–16.

15 Henrich K-O, Bauer T, Schulte J, et al. CAMTA1, a 1p36 Tumor Suppressor Candidate, Inhibits Growth and Activates Differentiation Programs in Neuroblastoma Cells. Cancer Res. 2011;71(8):3142–3151.

16 Alves G, Ornellas MH, Liehr, T. The role of calmodulin Binding Transcription Activator 1 (CAMTA1) gene and its putative genetic partners in the human nervous system. Psychogeriatrics. 2022;22:869–878.

17 Jiao S, Guan J, Chen M, et al. Targeting IRF3 as a YAP agonist therapy against gastric câncer. J. Exp. Med. 2018;215(2):699–718.

18 Li P, Ge D, Li P, et al. CXXC finger protein 4 inhibits the CDK18-ERK1/2 axis to suppress the immune escape of gastric cancer cells with involvement of ELK1/MIR100HG pathway. Cell Mol Med. 2020;24:10151–10165.

19 Kim H, Feng Y, Murad R, et al. Melanoma-intrinsic NR2F6 activity regulates antitumor Immunity. Sci. Adv. 2023;9:eadf6621.

20 Kavarthapu R, Morris C-HT, Dufau L. Prolactin induces up-regulation of its cognate receptor in breast cancer cells via transcriptional activation of its generic promoter by cross-talk between ERα and STAT5. Oncotarget. 2014;5(19):9079–9091.

21 Tang D, Kang R. NFE2L2 and ferroptosis resistance in cancer therapy. Cancer Drug Resist. 2024;7:41.

22 Li D, Lyu G. Prognostic Implications and Therapeutic Potential of MXD Genes in Gastric Cancer. Current Medicinal Chemistry. 2025;XX:1–20.

23 Hu C-L, Chen B-Y, Li Z, et al. Targeting UHRF1-SAP30-MXD4 axis for leukemia initiating cell eradication in myeloid leucemia. Cell Research. 2022;32:1105–1123.

24 Tagliabue E, Raimondi S, Gandini S. Meta-analysis of vitamin D–binding protein and cancer risk. Cancer Epidemiol Biomarkers Prev. 2015; 24(11):1758–1765.

25 Ushijima K, Suzuki C, Kitamura H, et al. Expression of clock gene Dbp in omental and mesenteric adipose tissue in patients with type 2 diabetes. BMJ Open Diab Res Care. 2020;8:e001465.

26 Chen X, Kang R, Kroemer G. Broadening horizons: the role of ferroptosis in cancer, Nat. Rev. Clin. Oncol. 2021;18:280–296.

27 Peng K, Ren X, Ren Q. NcRNA-mediated upregulation of CAMK2N1 is associated with poor prognosis and tumor immune infiltration of gastric cancer. Frontiers in Genetics. 2022;13:1–16.

28 Mohamed AA, Esmail OE, Ibrahim AMA, et al. The role of PRDM1 gene polymorphism in the progression of hepatocellular carcinoma in Egyptian patients. J Med Virol. 2023;95(1):e28343.

29 Li Q, Zhang L, You W, et al. PRDM1/BLIMP1 induces cancer immune evasion by modulating the USP22-SPI1-PD-L1 axis in hepatocellular carcinoma cells. Nature Communications. 2022;13(7677):1–17.

30 Jiang Y, He R, Jiang Y, et al. Transcription factor NFAT5 contributes to the glycolytic phenotype rewiring and pancreatic cancer progression via transcription of PGK1. Cell Death and Disease. 2019;10:948.

31 Zhao E, Yang S, Wen. Expression of NFAT5 gene in esophageal cancer tissues and its effect on the migration ability of esophageal cancer cells. Chinese Journal of Endocrine Surgery. 2023;6:681–685.

32 Chen X, Wang W, Li Y, et al. MYSM1 inhibits human colorectal cancer tumorigenesis by activating miR-200 family members/CDH1 and blocking PI3K/AKT signaling. J Exp Clin Cancer Res. 2021;40:341.

33 Garry DJ, Maeng G, Garry MG. Foxk1 regulates cancer progression. Ann Transl Med. 2020;8(17):1041.

34 Choi EJ, Seo EJ, Kim DK, et al. FOXP1 functions as an oncogene in promoting cancer stem cell-like characteristics in ovarian cancer cells. Oncotarget. 2015;7(3):3506–3519.

35 Iwafuchi M, Cuesta I, Donahue G, et al. Gene network transitions in embryos depend upon interactions between a pioneer transcription factor and core histones. Nat Genet. 2020;52(4):418–427.

36 Paris A, Tardif N, Galibert M.-D, Corre S. AhR and Cancer: From Gene Profiling to Targeted Therapy. Int. J. Mol. Sci. 2021;22:752.

37 Yan J, Tsai SY, Tsai M-J. SRC-3/AIB1: transcriptional coactivator in oncogenesis. Acta Pharmacologica Sinica. 2006;27(4):387–394.

38 Zhang Y, Wang B, Song H, Han M. GLIS3, a novel prognostic indicator of gastric adenocarcinoma, contributes to the malignant biological behaviors of tumor cells via modulating TGF-β1/TGFβR1/Smad1/5 signaling pathway. Cytokine. 2023;170:156342.

39 Ding Y, Wang Z, Chen C, et al. The gene regulatory molecule GLIS3 in gastric cancer as a prognostic marker and be involved in the imune infiltration mechanism. Front. Oncol. 2023;13:1091733.

40 Zhang Y, Wang X, Liu W, et al. CircGLIS3 promotes gastric cancer progression by regulating the miR⍰1343⍰3p/PGK1 pathway and inhibiting vimentin Phosphorylation. Journal of Translational Medicine. 2024;22:25.

41 Quiat D, Timberlake AT, Curran JJ, et al. Damaging variants in FOXI3 cause microtia and craniofacial microsomia. Genet Med. 2023;25(1):143–150.

42 Mao K, Borel C, Ansar M, et al. FOXI3 pathogenic variants cause one form of craniofacial microsomia. Nature Communications. 2023;14(2026):1–16.

43 Fan W, Cao DY, Yang B, et al. Hepatic Prohibitin 1 and Methionine Adenosyltransferase α1 Defend Against Primary and Secondary Liver Cancer Metastasis. J Hepatol. 2024;80(3):443–453.

44 Zhang H, Zhang G, Xiao M, et al. Two-polarized roles of transcription factor FOSB in lung cancer progression and prognosis: dependent on p53 status. Journal of Experimental & Clinical Cancer Research. 2024;43:237.

45 Gao Q, Wu Y, Lu C, et al. Knockdown of RREB1 inhibits cell proliferation via enhanced p16 expression in gastric câncer. Cell Cycle. 2021;20(23):2465– 2475.

46 Li T, Bou-Dargham M, Fultang JN, et al. c-Rel-dependent monocytes are potent immune suppressor cells in câncer. J Leukoc Biol. 2022;112(4):845–859.

47 Rahrmann EP, Wolf NK, Otto GM, et al. Sleeping beauty screen identifies RREB1 and other genetic drivers in human B-cell lymphoma. Mol Cancer Res. 2019;17(2):567–582.

48 Kabacaoglu D, Ruess DA, Ai J, et al. NF-κB/Rel Transcription Factors in Pancreatic Cancer: Focusing on RelA, c-Rel, and RelB. Cancers. 2019;11:937.

49 Takahashi H, Varner J. Rel-ating myeloid cells to cancer therapy. Nature Cancer. 2020;1:480–481.

50 Cloud AS, Vargheese AM, Gunewardena S, et al. Loss of REST in breast cancer promotes tumor progression through estrogen sensitization, MMP24 and CEMIP overexpression. BMC Cancer. 2022;22:180.

51 Redin E, Sridhar H, Zhan YA, et al. SMARCA4 controls state plasticity in small cell lung cancer through regulation of neuroendocrine transcription factors and rest splicing. Journal Of Hematology & Oncology. 2024;17:58.

52 Wang J, Xiao K, Hou F, et al. POU2F1 Promotes Cell Viability and Tumor Growth in Gastric Cancer through Transcriptional Activation of lncRNA TTC3-AS1. Journal of Oncology. 2021;2021(5570088):15.

53 Wang J, Hou F, Tang L, et al. The interaction between long non⍰coding RNA LINC01564 and POU2F1 promotes the proliferation and metastasis of gastric cancer. Journal of Translational Medicine. 2022;20:220.

54 Xiao Y, Yang P, Xiao W, et al. POU2F1 inhibits miR-29b1/a cluster-mediated suppression of PIK3R1 and PIK3R3 expression to regulate gastric cancer cell invasion and migration. Chinese Medical Journal. 2025;138(7):838–850.

